# STRESS, an automated geometrical characterization of deformable particles for *in vivo* measurements of cell and tissue mechanical stresses

**DOI:** 10.1101/2021.03.26.437148

**Authors:** Ben Gross, Elijah Shelton, Carlos Gomez, Otger Campàs

## Abstract

From cellular mechanotransduction to the formation of embryonic tissues and organs, mechanics has been shown to play an important role in the control of cell behavior and embryonic development. Most of our existing knowledge of how mechanics affects cell behavior comes from *in vitro* studies, mainly because measuring cell and tissue mechanics in 3D multicellular systems, and especially *in vivo*, remains challenging. Oil microdroplet sensors, and more recently gel microbeads, use surface deformations to directly quantify mechanical stresses within developing tissues, *in vivo* and *in situ*, as well as in 3D *in vitro* systems like organoids or multicellular spheroids. However, an automated analysis software able to quantify the spatiotemporal evolution of stresses and their characteristics from particle deformations is lacking. Here we develop STRESS (Surface Topography Reconstruction for Evaluation of Spatiotemporal Stresses), an analysis software to quantify the geometry of deformable particles of spherical topology, such as microdroplets or gel microbeads, that enables the automatic quantification of the temporal evolution of stresses in the system and the spatiotemporal features of stress inhomogeneities in the tissue. As a test case, we apply these new code to measure the temporal evolution of mechanical stresses using oil microdroplets in developing zebrafish tissues. Starting from a 3D timelapse of a droplet, the software automatically calculates the statistics of local anisotropic stresses, decouples the deformation modes associated with tissue- and cell-scale stresses, obtains their spatial features on the droplet surface and analyzes their spatiotemporal variations using spatial and temporal stress autocorrelations. The automated nature of the analysis will help users obtain quantitative information about mechanical stresses in a wide range of 3D multicellular systems, from developing embryos or tissue explants to organoids.

**Author summary:** The measurement of mechanical stresses in 3D multicellular systems, such as living tissues, has been very challenging because of a lack in technologies for this purpose. Novel microdroplet techniques enable direct, quantitative *in situ* measurements of mechanical stresses in these systems. However, computational tools to obtain mechanical stresses from 3D images of microdroplets in an automated and accurate manner are lacking. Here we develop STRESS, an automated analysis software to analyze the spatiotemporal characteristics of mechanical stresses from microdroplet deformations in a wide range of systems, from living embryonic tissues and tissue explants to organoids and multicellular spheroids.

## Introduction

Mechanical forces are known to play an essential role in the control of cell behavior as well as in tissue and organ morphogenesis. During embryogenesis, cells apply differential forces to guide tissue flows and build embryonic structures [1, 2]. While mechanical forces are generated at the cell and subcellular scales, it is the collective force generation by many cells and its transmission over supracellular length scales that shapes functional tissues [3]. Understanding the characteristics of mechanical stresses at different length- and time-scales is necessary to bridge the gap between force generation at the cell scale and organ formation at much larger scales [1–3]. Beyond their important role in morphogenesis, mechanical forces are also known to play an important role in the control of cell behavior. Mechanobiology studies have shown that important cell behaviors, such as cell differentiation or proliferation, are affected by the mechanical forces acting on them [4, 5]. Therefore, having the ability to quantify the spatiotemporal characteristics of mechanical stresses in 3D multicellular systems can help advance our understanding of how mechanics affects both individual cell behavior and their collective dynamics.

While several measurement techniques exist to quantify forces in 2D cell monolayers on synthetic culture dishes [6], techniques to measure mechanical forces in 3D multicellular environments are scarce [7, 8]. The development of microdroplet techniques enabled the quantification of forces in 3D multicellular environments, from 3D cell culture to developing tissues, *in vivo* and *in situ* [9–12]. Mirroring oil microdroplets, gel microbeads have been recently used to measure stresses in multicellular systems [13–16] and also for single cell studies [17]. Both oil microdroplets and gel microbeads enable measurements of mechanical stresses in various 3D systems, allowing quantitative mechanobiology studies in a more physiological context.

The accurate measurement of mechanical stresses with deformable particles relies on precise measurements of surface deformations of the probe used (oil microdroplets or gel microbeads). Oil microdroplets enable precise measurements of anisotropic stresses solely from knowledge of their surface geometry and the droplet interfacial tension (which can be calibrated *in vivo* and *in situ*) [9, 12], whereas measurements of stresses with gel microbeads require the quantification of strain fields inside the probe [15]. However, approximate methods have been recently developed to obtain mechanical stresses using gel microbeads from their surface deformations and the bead stiffness [17]. While different analysis methods exist to reconstruct the geometry of the probes in 3D [12, 17, 18], an automated and and reliable tool to obtain stresses in 3D and time, as well as to analyze their spatiotemporal characteristics, is lacking.

Taking advantage of methods from computational differential geometry, we developed STRESS (Surface Topography Reconstruction for Evaluation of Spatiotemporal Stresses), a fully automated code for surface reconstruction and analysis that enables the calculation of multiple geometric aspects of the probe surface, ranging from its curvatures, to surface integrals and geodesic distances. We use the extensive characterization of surface geometry to characterize multiple aspects of the measured stresses. The complete pipeline starts from a 3D timelapse of a deformable particle (in the test case used here, this is an oil microdroplet inserted in developing zebrafish embryo) and automatically reconstructs the surface deformations of the particle, obtains its geometrical characteristics and analyzes the spatiotemporal changes of mechanical stresses. STRESS automatically decouples mechanical stresses occurring at supracellular (tissue) scales and cell-scale by analyzing different deformation modes as well as maximal and minimal values of stress on the droplet surface. It also obtains the characteristics of spatial inhomogeneities of the stresses around the droplet, providing insight on the spatial structure of the stresses in the tissue. Moreover, it automatically calculates the persistence of total stresses, as well as of cell- and tissue-scale stresses, from temporal autocorrelation functions. Finally, the analysis also provides accurate measurements of the probe volume over time, which facilitate the measurement of isotropic stresses (tissue pressure) using gel microbeads. An open source version of STRESS, with specific instructions on how to install it and use it, can be downloaded from https://github.com/campaslab/STRESS.

## Materials and methods

### Imaging and sample preparation

Imaging and sample preparation of surface-labeled microdroplets in tooth mesenchymal cell aggregates and volume-labeled (or interior-labeled) droplets in developing zebrafish tissues was done as previously described in references [12] and [9], respectively. Surface-labeled droplets were coated with Cy5–streptavidin. Volume-labeled droplets were prepared by dissolving FCy5 dye [19] in Novec 7300 fluorocarbon oil (3M) at a 25 *μ*M concentration. A fluorinated Krytox-PEG(600) surfactant (008-fluorosurfactant, RAN Biotechnologies) was also diluted in the fluorocarbon oil at a 2.5% (w/w) concentration. This fluorescently-labeled oil was injected directly in the embryo, as previously described [9], and the volume of the droplet was set by the injection pressure and the duration of the injection.

### Obtaining the point cloud

A point cloud representation of the droplet surface is constructed by tracing rays (or lines) through the droplet surface in the fluorescence 3D image of the droplet and fitting the sampled intensity profiles with an appropriate edge profile model, which depends on what feature of the droplet is labelled fluorescently. Ray tracing paths were determined as previously described [18]. For droplets labeled at their surface (Fig. 1a), we employ a Gaussian edge profile model [18] (Fig. 1d). However, for droplets prepared with fluorescently labelled fluorocarbon oil [9, 19, 20] (Fig. 1b), which display fluorescence signal throughout the droplet, we used an attenuated sigmoid profile model that captures both the fluorescence intensity fast change at the droplet surface and the mild attenuation of fluorescence inside the droplet (due to imaging a 3D object). Specifically, we use the fit model *F* (*x*) = (*d* + *c* * (*x* − *b*))/(1 + exp(*a* * (*x* − *b*))) + *e* for the intensity profile along the traced rays within the image volume at distance *x* from a starting point 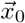 along the ray direction 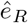 (oriented in the direction of the outward surface normal vector). Parameters *a, b, c, d* and *e* are obtained from nonlinear least squares fitting using the Levenberg-Marquardt algorithm implemented with the Matlab (MathWorks) function lsqnonlin(). The location of the surface, which corresponds to a single point 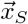 along the ray, is determined from *b*, namely 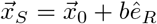.

**Fig 1.**
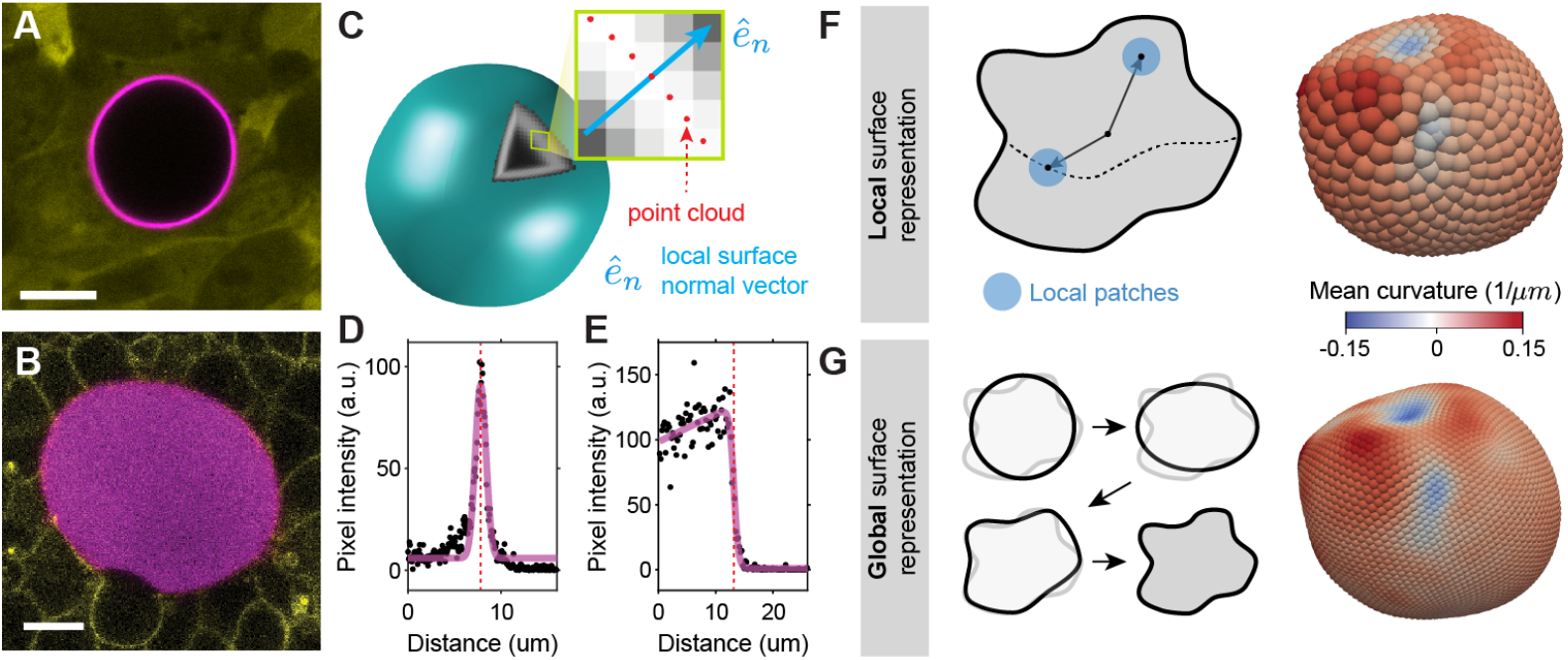
Particle’s surface reconstruction from 3D fluorescence microscopy images. **A-B**, Examples of confocal sections of microdroplets fluorescently labelled at their surface (**A**; magenta) and in their interior (**B**; magenta). Before imaging, the droplets were inserted in a 3D aggregate of tooth mesenchymal cells (**A**; cytoplasmic label, yellow) and in the presomitic mesoderm of developing zebrafish embryos (**B**; cell membranes, yellow). **C**, Sketch of a deformed particle labelled fluorescently at its surface. The inset shows a detail of the particle surface with its local normal vector 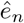 and the detected surface points (point cloud; red). **D-E**, Fluorescence intensity along rays traced along the normal direction to the surface for both surface labelled (**D**) and internally labelled droplets (**E**). Fits to the intensity profile along a ray (magenta) are used to detect the location of the surface (red dashed line). **F**, A local representation of the surface geometry can be done using Monge patches to estimate the local mean curvature at every point on the droplet surface. **G**, A global representation of the surface geometry can be obtained using a spherical harmonic basis, which progressively capture finer details as the number of modes is increased. The contribution of each mode depends on every point on the surface. The mean curvature of the deformed particle can be obtained at every point of the surface with high spatial resolution.

### Global surface representation

Once a point cloud representing the droplet surface has been obtained, we use a spherical harmonic basis to obtain a global approximation of the surface. A global function approximating the object surface is constructed by finding the least-squares fit of spherical harmonics to the point cloud [21], up to a given degree. The degree of the fit is limited by the number of points in the point cloud, since we need at least one degree of freedom in our data for each degree of freedom in our fit. However, in order to have the fitting functions properly constrained, we restrict ourselves to two degrees of freedom (data points) in our point cloud for each degree of freedom in our fit. This means that we would ideally fit no more than 50 global basis functions to a point cloud with 100 points. The spherical harmonic fits provide a smooth representation of the surface with spherical topology, whose variations are limited by the maximum degree of harmonics chosen. Importantly, the spherical harmonics converge spectrally to smooth functions on the sphere [21], so it is possible to achieve a faithful representation of a smooth droplet surface with a small to moderate number of modes. This approach, adapted from recent work on surface flows on thin-shell membranes [22–24], has also been used recently to analyze cellular shapes [25].

### Mean curvature map

Once we have a smooth global representation of the droplet surface, we calculate its mean curvature, which depends on the coordinate system chosen to parameterize our global fit. In the simplest case, we define the surface points in a cartesian coordinate system, namely **r_s_** = (*x_s_*(*θ, ϕ*), *y_s_*(*θ, ϕ*), *z_s_*(*θ, ϕ*)), where the coordinates of each point are given in terms of the azimuthal and polar coordinates, *θ* and *ϕ* respectively, of our spherical harmonic basis. By taking the first and second derivatives of our spherical harmonic basis functions, we can calculate any geometric quantity associated to the droplet surface, including the mean curvature *H* and the Gaussian curvature *K*, using standard differential geometry [26].

All of the calculations are done using python’s numpy library [27]. We validated the convergence of the representations of these geometric quantities using the method of manufactured solutions [23, 24]. The symbolic expressions needed for these validation tests, performed on various surface geometries, were generated by python’s sympy package [28].

### Surface integrals and particle volume

One major advantage of using spherical harmonics to obtain a global surface representation is that, for a given degree, there is a set of lebedev quadrature points that integrates these harmonics exactly on the surface of the unit sphere [29]. For our calculations, we use a quadrature of up to 5810 lebedev points, which can integrate inner-products of up to degree 65 spherical harmonics exactly. We specifically use these lebedev points to represent the surface in our global fit and also to display the surfaces.

In order to calculate the object’s (microdroplet or gel microparticle) volume *V*, we adapt our surface quadrature. Parameterizing the droplet surface in spherical coordinates (*r*, *θ*, *ϕ*) the object’s volume is given by

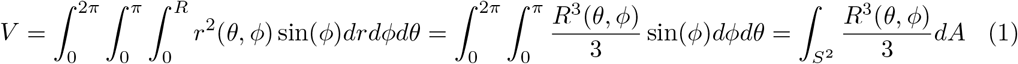

where *R*(*θ, ϕ*) is the droplet radius at each point of the surface and *S*^2^ denotes the surface of the unit-sphere. We tested the accuracy of volume measurements by calculating the volume of a very eccentric ellipsoid (with semi-axes lengths 3 *μ*m, 5 *μ*m, and 7 *μ*m) using Eq. 1 and comparing it to its analytically known volume of 140 *πμ*m^3^. Using only 590 quadrature points (much less than the 5810 quadrature points used for all our calculations), we find the relative error in the ellipsoid volume to be ≃ 10^−5^%.

### Tissue-scale stresses

To obtain the ellipsoidal mode of the droplet shape, we perform a least-squares ellipsoidal fit on the segmented point cloud. Specifically, we fit the equation: *Ax*^2^ + *Bxy* + *Cy*^2^ + *Dxz* + *Eyz* + *Fz*^2^ + *Gx* + *Hy* + *Iz* = 1 to the point cloud and to find the 9 coefficients that represent the least squares ellipsoid. To obtain the mean curvature of this ellipsoid, we transform this ellipsoid into the coordinates system (*x*_1_, *x*_2_, *x*_3_) defined by its principal directions. These coordinates are along the major, medial, and minor axes of the ellipsoid, respectively, with the same center, namely

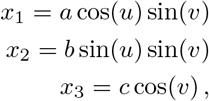

where *u* ∈ [0, 2*π*) and *v* = [0, *π*] parameterize the ellipsoid surface. From this parameterization, we can use a closed form expression [30] to calculate the mean curvature of the ellipsoid, *H_e_*, at every point of the surface, which reads

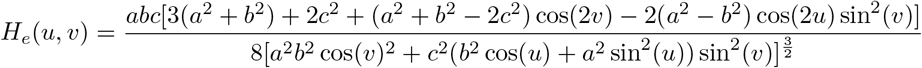

Using this expression, it is possible to obtain a closed form expression for the anisotropic tissue stresses 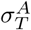. The maximum mean curvature 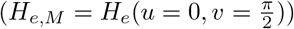 is *H_e,M_* = *a*/(2*c*^2^) + *a*/(2*b*^2^) and the minimum mean curvature (*H_e,N_* = *H_e_*(*u* = 0, *v* = 0)) is *H_e,N_* = *c/*(2*b*^2^) + *c*/(2*a*^2^). The anisotropic tissue stresses 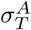 are given by the maximum stress anisotropy between two principal directions and is directly related to the maximal and minimal mean curvatures, namely

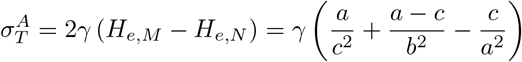

where *γ* is the droplet interfacial tension.

### Cell-scale stresses

To quantify the stresses arising from higher order deformation modes (beyond the ellipsoidal mode), which are associated with length scales closer to the cell size, we calculate the deviations of the droplet deformations from the ellipsoidal mode. For any given timepoint, we subtract the calculated total anisotropic stress value *σ^A^* from the tissue-scale stresses associated with the ellipsoidal deformation, namely 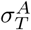, to obtain the cell-scale stresses (Eq. 6). Specifically, to compute the cell-scale stress anisotropy between points *p* and *q* on the surface, we compute 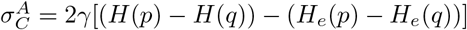, where we choose the points *p* and *q* on the ellipsoid that correspond with the same ellipsoidal coordinates as *p* and *q* on the droplet.

In order to obtain the extrema on the deformed surface, we use the surface triangulation of the 5810 lebedev nodes on the surface to relate them to each other. Two points are classified as neighbors if they are connected by an edge. Local maxima are points whose value of cellular stress is greater or equal to each of their neighbors’ values, and local minima are points whose value of cellular stress is less than or equal to each of their neighbors’ values. The extrema stresses (or adjacent extrema stresses) are obtained by calculating the difference in cell-scale stresses between maxima and minima (or adjacent maxima and minima).

### Geodesic distances

To calculate distances between points on the curved droplet surface, we first triangulate the surface using a Delauney Triangulation of the lebedev quadrature points on the sphere using python’s scipy package [31]. The geodesic distance between lebedev quadrature points on the droplet surface is calculated from the surface triangulation graph using using the gdist package [32]. To obtain maximal resolution and minimize noise, we use the full set of 5810 lebedev nodes in our calculations.

### Autocorrelation functions

To obtain the spatial autocorrelation of stresses on the droplet surface at a given timepoint *t_i_*, we compute the product of each pair of stress values separated by a geodesic distance *s* between them. To bin the points, we use a bump function *w*(*s*) = exp 1 − 1/(*ℓ*^2^ − *s*^2^) for *s* < *ℓ*, where we pick *ℓ* to be 5% of the geodesic diameter of the droplet. To calculate the spatial autocorrelations we incorporate this weight in Eq. 7

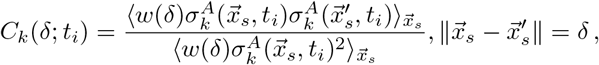

where *k* indicates that this calculation can be performed for any stress field on the droplet surface, including the total stress *σ^A^* and the tissue- and cell-scale stresses 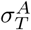 and 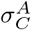, respectively. We have checked that changing the value of *ℓ* does not change our results. Very small values of *ℓ* (less than 1% of the geodesic diameter of the droplet) lead to more noisy results for the autocorrelation, but do not change the features of the obtained autocorrelation function.

For temporal correlations, we use a radial parameterization of the droplet (a system of spherical coordinates with the same center as the droplet). This allows us to relate points on the droplet surface with the same spherical coordinates as the droplet changes over time. We then calculate the temporal correlations of the stress values at the same spatial location (same angular coordinates) and average the results for all the points on the droplet surface (Eq. 8).

## Results

To explain and exemplify the STRESS analysis pipeline, we focus on the use of microdroplets to measure mechanical stresses in living tissues, as in this case measurements of stresses can be obtained solely from the droplet surface deformations (knowing the droplet interfacial tension). We performed 3D timelapse imaging of an oil microdroplet in the presomitic mesoderm of a zebrafish embryo for 60 minutes (Methods), and use STRESS to characterize spatiotemporal features of the measured stresses.

### Geometrical representation of the particle’s surface

In order to measure stresses from deformable particles and, in particular, from oil microdroplets, it is necessary to image the particles in 3D. While it is also possible to measure stresses from partial 3D microdroplet reconstructions and even from 2D confocal sections [9, 12], the analysis presented herein focuses on complete particle reconstructions. For measurements in 3D multicellular systems, deformable particles are first inserted between the cells of developing embryos or multicellular aggregates [9, 10, 12–16], whereas for experiments with single cells in culture the particles are simply brought in contact with the cells [12, 17]. Typically, particles are fluorescently labeled at their surface (Fig. 1A) and/or interior (Fig. 1B), allowing their 3D imaging with various microscopy techniques, such as confocal or lightsheet microscopy. Once the 3D image of the droplet is obtained, the location of its surface needs to be detected to obtain its mathematical representation.

Previous raytracing algorithms have been shown to provide a faithful representation of the particle’s surface for surface-labeled deformable particles [18]. This method identifies first the center of the droplet and traces rays in all spatial directions (Fig. 1C). By measuring the intensity profile along the rays and detecting the intensity maximum, it is possible to detect the location of the surface along the ray with high spatial resolution (Fig. 1D). Repeating this procedure for all the rays provides a point cloud that represents the surface of the object. A refined point cloud is then obtained by repeating the procedure for rays retraced along normal directions to the surface [18] (Fig. 1C). We have extended this algorithm for particles fluorescently labeled in their interior (Fig. 1B). To do so, we employ the same raytracing procedure, but implement a different fitting function for the intensity profile along the traced rays because particles with interior label display a high fluorescence signal in their interior and a sharp decrease at their surface (Fig. 1B,E; Methods). Fitting the intensity profile along a ray allows the detection of the surface location on that same ray (Fig. 1E; Methods). By repeating this procedure along all traced rays, we obtain the point cloud that represents the location of the particle’s surface.

Once the point cloud that represents the deformed particle’s surface has been determined, there are several approaches to characterize its geometry. We previously developed a methodology to obtain the surface mean curvature map from local quadratic fits [18] (Fig. 1F). This strategy relies on the construction of local Monge patches of adaptable size and the approximation of the local surface geometry within the patch. Because this method employs only on local information, it allows the calculation of curvature maps on both fully and partially reconstructed particles, but is more prone to errors associated with limited spatial resolution or poor signal-to-noise ratios, especially for very small patch sizes. To overcome these issues in full 3D particle reconstructions, we mathematically characterize the surface using global fits of controlled degree (Fig. 1G). Specifically, we use least-squares fits of spherical harmonics (up to a desired degree) to the segmented point cloud (Methods). The obtained surface representation using global fits provides a smooth, highly-resolved and accurate mathematical representation of the surface that is less sensitive to high frequency noise in the point cloud, as each fits contains information from all points on the surface (Fig. 1G). This global surface representation can be used to quantify a wide range of geometrical features, from mean and gaussian curvature maps to geodesic distances, with high spatial resolution and accuracy. In particular, STRESS calculates the mean curvature 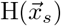 at every point 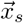 on the droplet surface, as this geometrical quantity is central to the calculation of stresses using deformable particles.

### Quantifying the temporal evolution of anisotropic stresses from droplet deformations

As a test case for STRESS, we measure the time evolution of local stresses in 3D developing zebrafish tissues by performing 3D timelapses of volume-labeled microdroplets that were previously inserted in the tissue (Methods). For each timepoint *t_i_* (*i* = 1, …, *N*, with *N* being the number of frames in the 3D timelapse), we reconstruct the droplet in 3D, calculate its point cloud, and obtain its 3D smooth surface representation as well as the mean curvature map 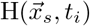, as described above. This procedure results in the time evolving mean curvature map of the droplet surface, 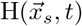. As described in the remaining part of the manuscript, the droplet deformations encode different stresses and features of the spatiotemporal evolution of the stresses in the tissue. We first define the total anisotropic stresses, 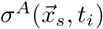, which, for oil microdroplets, are obtained directly from the mean curvature map as previously described [12], namely

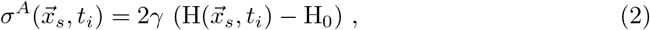

where *γ* is the droplet interfacial tension and H_0_ is the average of the mean curvature on the droplet surface. The measured total anisotropic stresses are the stresses responsible for deforming the droplet from the spherical geometry and are caused by mechanical stress inhomogeneities surrounding the droplet. There are multiple ways to measure the interfacial tension to calibrate the microdroplets, as previously described [9, 11, 12]. For the specific example here, we measured the interfacial tension *in vivo* and *in situ* using magnetically-responsive microdroplets (*γ* ≃ 3.3 mN/m; Methods). Knowing the droplet interfacial tension, we use Eq. 2 to obtain the 3D map of total anisotropic stresses on the droplet surface at each timepoint (Fig. 2A,B), as the droplet follows morphogenetic flows in the tissue. Since the global surface representation of the droplet enables the accurate calculation of integrals on the surface (Methods), we calculate the average mean curvature H_0_ as H_0_ = (*∫*_*M*_ *HdA*)/(*∫*_*M*_ *dA*), where *M* is the (manifold) droplet surface (Fig. 2C; Methods). Directly averaging the mean curvature values does not provide an accurate measure of the average mean curvature (Fig. 2C). For this reason, all our calculations obtain the value of the average mean curvature from its surface integral.

**Fig 2.**
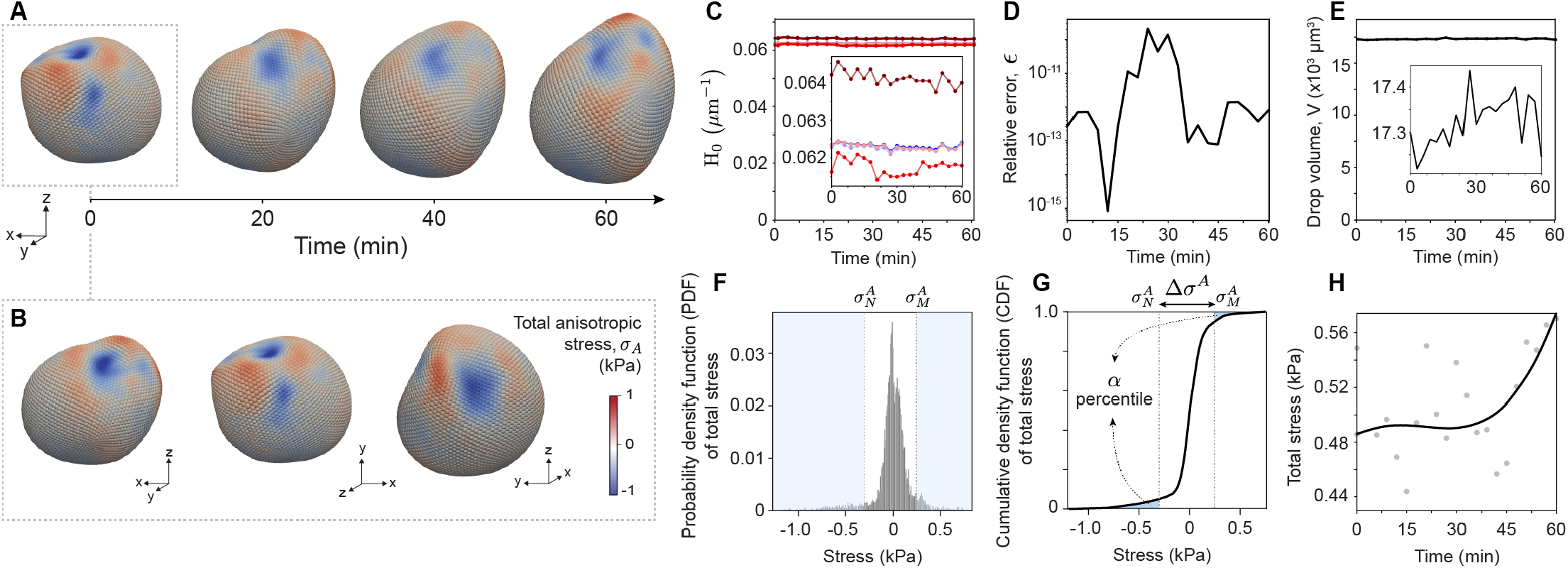
Quantifying temporal changes in anisotropic stresses using oil microdroplets. **A**, Snapshots of the total anisotropic stresses mapped on the droplet surface at different timepoints. **B**, Different perspectives of the stresses on the droplet surface at the initial timepoint. **C**, Time evolution of the average mean curvature *H*_0_ using distinct methodologies. Accurate results are obtained by computing the surface integral of the mean curvature map (light blue), by integrating the droplet volume to obtain the undeformed droplet radius (dark blue) and by calculating the surface integral of the mean curvature map of the ellipsoidal deformation mode of the droplet (pink; Methods). Values of the average mean curvature obtained by performing ensemble averages of the mean curvature values at segmented points (dark red) or at lebedev points (light red) deviate from the properly calculated average obtained using the surface integral of the mean curvatures. **D**, Relative error of the calculation, obtained using the Gauss-Bonnet theorem. This metric allows us to detect inconsistencies in the droplet surface due to poor segmentation. **E**, Calculated time evolution of the droplet volume, *V*. Small fluctuations in the volume (~ 1%) due to imaging noise are observed. **F-G**, Probability distribution function (or normalized frequency; **F**) and cumulative distribution function (**G**) of total anisotropic stresses, *σ^A^*, on all 5810 lebedev points. The 5^*th*^ and 95^*th*^ percentile values (*α* = 0.05) define the minimal and maximal anisotropic stresses, 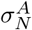 and 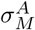 respectively. **h**, Time evolution of the magnitude of total anisotropic stresses, Δ*σ^A^*. B-spline (black line) shows the trend and gray points are the measured values.

In order to evaluate the accuracy of the global surface representation, we make use of the Gauss-Bonnet Theorem, which states that the integral of the Gaussian curvature *K* over any smooth surface with spherical topology should equal 4*π*. We integrate the Gaussian curvature *K* over the droplet surface and measure the relative deviation (or error), *∊*, from the expected 4*π*, namely *∊* ≡ |1 − *∫*_*M*_ *KdA*/4*π*, at each timepoint (Fig. 2D). Inaccurate droplet segmentations (i.e., those containing errant points or large holes in the point cloud) will cause the error *\** to increase as the degree of the surface representation increases, allowing the automated detection of incorrectly segmented images, which we do not include in our measurements (Methods). For the example shown here, the relative errors are very small (Fig. 2D), indicating no issues with droplet segmentation, and fluctuate over time as the droplet segmentation is slightly different at each timepoint.

In addition to surface integrals, it is also possible to accurately calculate volume integrals using the global representation of the particle’s surface, enabling the monitoring of the particle’s volume *V* over time (Methods). While oil droplets are incompressible and no volume changes are expected over time, certain oils may have a small but finite solubility in aqueous media, which could cause the droplet volume to decrease over time. For the specific oil and conditions in our experiments (Methods), we did not observe any significant reduction in droplet volume (Fig. 2E). The observed volume fluctuations are of approximately of 1.6% and due to imaging noise, as the accuracy in the volume calculation is much higher (Methods). It is also possible to use the droplet volume to accurately calculate the average value of the mean curvature of the droplet, namely 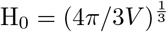 (Fig. 2D). In contrast to microdroplets, changes in the volume of gel microbeads provide a measure of the local isotropic stress or tissue pressure. Therefore, the ability to accurately measure volume over time can be employed to track temporal changes in tissue pressure when using gel microbeads.

While the full stress map on the droplet surface provides spatiotemporal information about stresses (Fig. 2A,B), in most applications it is necessary to define a single numerical value of the total anisotropic stress for a droplet at a given timepoint, so that it is possible to perform statistics and monitor the temporal evolution of stresses in the tissue. To do so, we first calculate the statistical distribution of the magnitude of total anisotropic stresses from all the values of the stresses at different points on the droplet surface (probability density function; Fig. 2F). This distribution has important information about the variations in the magnitude of local stresses in the tissue. We then obtain the cumulative distribution of anisotropic stresses (cumulative density function; Fig. 2G), and define the range of values at each extreme of the distribution accounting for an *α* percent of the data, thereby defining minimum and maximum stress values, 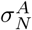 and 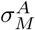 respectively. The amplitude of total anisotropic stresses between this maximum and minimum defines a single numerical measure of the total anisotropic stresses, namely 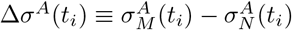. If *α* = 0, the minimal and maximal values correspond to the actual minimal and maximal values of the total anisotropic stress on the droplet surface. However, since extreme values are prone to noise, it is preferable to define a small value of *α* to cut off extreme fluctuations. Here we used *α* = 0.05 (or 5%; removing the 5% largest and 5% smallest values of the total stress). By measuring the magnitude of total anisotropic stresses at each timepoint, we can track the time evolution of total anisotropic stresses Δ*σ^A^*(*t*) (Fig. 2H).

### Tissue-scale (supracellular) stresses

As shown above, total anisotropic stresses can be calculated at each timepoint and monitored over time. However, the droplet deformations encode different types of stresses that contain distinct information about the mechanical state of the tissue. While high order deformation modes (localized higher/lower curvature changes) provide information about specific length scales, the lowest order deformation mode, namely the ellipsoidal mode, provides information about the mechanical stress anisotropy occurring at the length scale of the droplet size (Fig. 3A,B). By making droplets bigger than the cell size (typically ~ 4 cell diameters; Fig. 3B), the ellipsoidal deformation mode enables the measurement of the local value of the anisotropic stress at supracellular scales (tissue-scale), 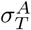. This tissue-scale stresses essentially average out smaller deformations on the droplet surface and are thus associated with stress anisotropy in the tissue propagating at length scales larger than the cell size.

**Fig 3.**
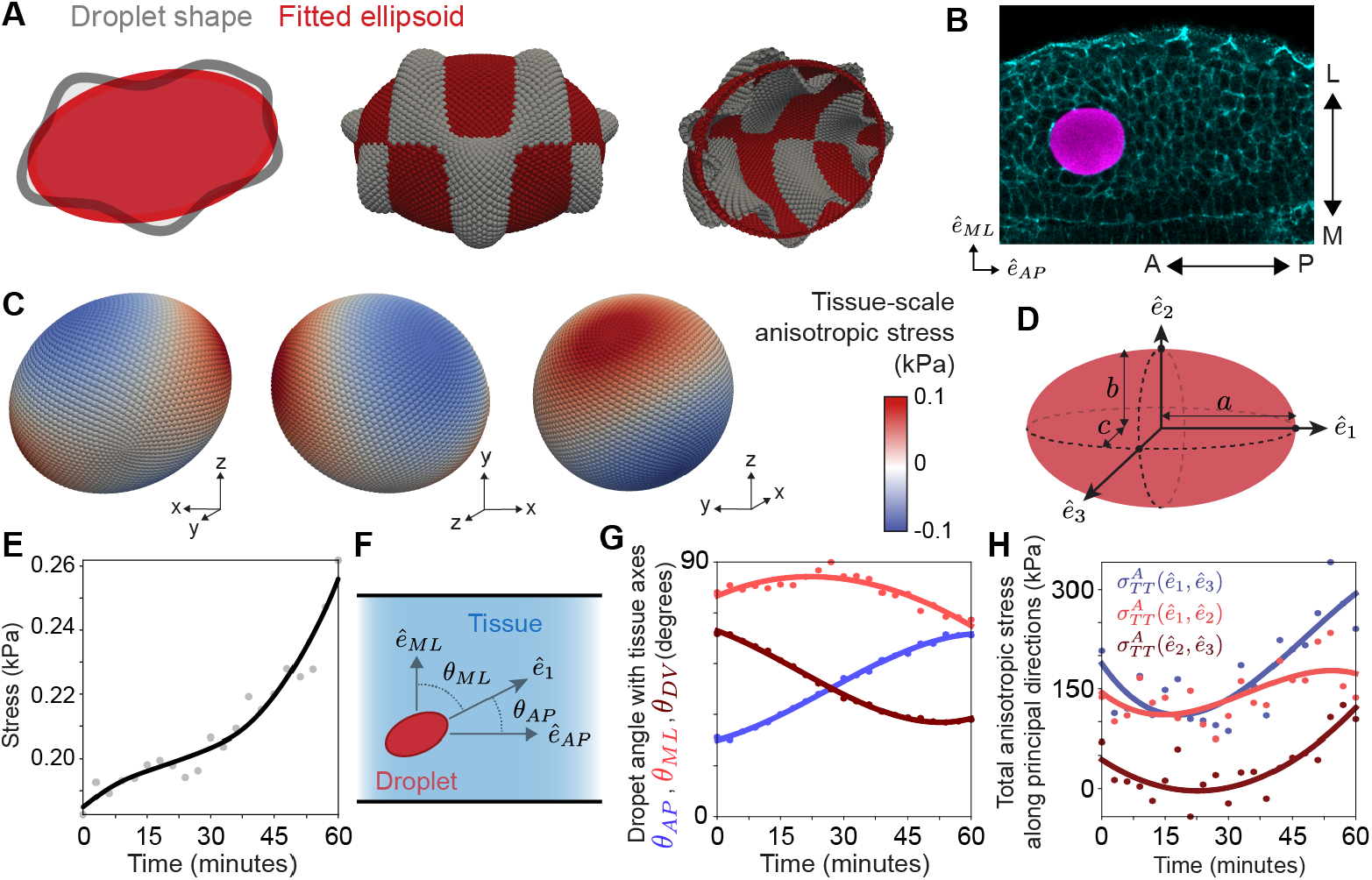
Quantifying tissue-scale (supracellular) stresses over time. **A**, Sketch of a 2D section (left) and of a deformed droplet (gray) and associated ellipsoidal fit (red). Cross-sections (center and right) of a 3D least-squares fit of an ellipsoid (red) to a manufactured dataset (grey). **B**, Confocal section of the pre-somitic mesoderm of a developing zebrafish embryo with an oil microdroplet in between the cells of the tissue (Methods). The relevant embryonic axes are defined: anteroposterior (AP) axis, mediolateral (ML) axis, and dorsoventral (DV) axis (DV axis not shown; perpendicular to the imaged embryo section). **C**, Tissue-scale stresses shown on the surface of the ellipsoidal droplet deformation mode at a fixed timepoint. **D**, Sketch of the ellipsoidal deformation mode (red), which is characterized in terms of the principal directions 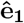, 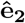 and 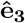, with semi-axes length *a*, *b* and *c*, respectively. **E**, Time evolution of tissue-scale stresses, 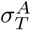. The b-spline show the trend and the gray points correspond to the measured values. **F**, Sketch of a droplet in the tissue defining the angles *θ_AP_*, *θ_ML_* and *θ_DV_* between the axis of droplet elongation, 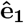, and the relevant embryonic axis, 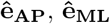 and 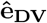, respectively. **G**, Time evolution of the ellipsoidal deformation orientation in the tissue. **H**, Time evolution of the total anisotropic stresses between principal directions.

We obtain the ellipsoidal droplet deformation by fitting an ellipsoid to the point cloud (Methods; Fig. 3C). The principal axes of ellipsoidal deformation, characterized by unit vectors 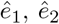 and 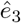 with corresponding semiaxes *a*, *b* and *c*, respectively (Fig. 3D), reveal the principal directions of stress anisotropy in the tissue. By construction, we choose *a* to be the longest semiaxis, and 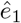 thus reveals the direction of droplet elongation. The tissue-scale stress anisotropy between two principal directions *i* and *j* (*i*, *j* = 1, 2, 3 and *i* ≠ *j*) is obtained by calculating the stress anisotropy between these directions, namely

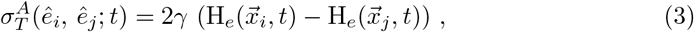

where H_*e*_ corresponds to the mean curvature of the ellipsoid and 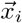 and 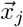 are the surface locations where the principal axis 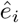 and 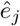 cross the ellipsoid surface (Fig. 3B). In order to obtain a single measure of the tissue-scale stress anisotropy, we calculate the difference between the maximal and minimal stresses on the ellipsoid (Methods), which are associated with two principal directions, namely

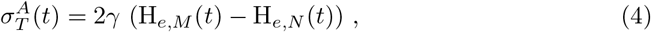

where H_*e,M*_ and H_*e,N*_ are the maximal and minimal mean curvature values of the ellipsoidal deformation mode. The measured tissue-scale anisotropic stresses display much smaller values than the total stress (Fig. 3E). This is because the tissue-scale stresses only account for the lowest deformation mode and higher deformation modes are associated with larger stresses. Unlike the total stresses, which remain roughly constant over time within the measurement error (Fig. 2H), the tissue-scale stresses display an increase of approximately 60% over 1 hour.

The principal directions reveal the local, tissue-scale mechanical anisotropy in the tissue. To quantify changes in the orientation of the mechanical stress anisotropy in the tissue, it is informative to monitor the time evolution of droplet orientation. To do so, it is necessary to define the biologically relevant spatial directions, or equivalently, the natural coordinate system of the tissue, which, in general, is different for distinct systems. For the example studied here, these directions are the anteroposterior axis 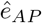, the mediolateral direction 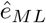 and the dorsoventral axis 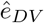 (Fig. 3B). The angle between 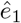, which quantifies the direction of droplet elongation, and each of these directions is given by 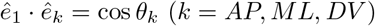 and their time evolution can be tracked by calculating these angles at each timepoint (Fig. 3F,G). The time evolution of these angles indicates that the droplet is consistently perpendicular to the ML axis for the time course of the experiment. However, the droplet rotates over time in the AP-DV plane, starting more aligned along the AP axis and progressively becoming more aligned along the DV axis, revealing a change in tissue-level stresses anisotropy in the AP-DV plane.

Beyond the tissue-scale stresses associated with the ellipsoidal droplet deformation, it is also possible to define the total stress anisotropy along the principal directions, namely

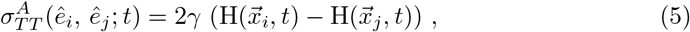

which differ from the the tissue-scale stresses defined above (Eq. 3) in that it employs the mean curvature H of the actual droplet surface at 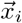 and 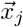 rather than the mean curvature of the ellipsoidal mode at these points. While the total stress anisotropy along principal directions 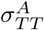 provides the actual value of stress anisotropy, it is more noisy than 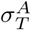 as it is affected by higher order deformations (Fig. 3H). In contrast, since 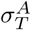 (Eq. 4) uses the mean curvature of the fitted ellipsoid, it removes the noise associated with higher order deformation modes and provides a better defined measure of tissue-scale stresses.

### Cell-scale stresses

To quantify cell-scale stresses we study the deviations of the droplet deformations from the ellipsoidal mode. We first express the droplet deformations on the ellipsoidal reference frame and then calculate the deviations of the deformations from the ellipsoidal mode (Fig. 4A), which, by definition, only contain deformation modes of higher order than the ellipsoidal one. To calculate the deviations from the ellipsoidal mode, we perform a least squares fit of the distances between the droplet surface and the ellipsoid, using the coordinate system of the fitted ellipsoid to parameterize the residuals (or deviations). For any timepoint *t_i_*, cell-scale stresses are given by

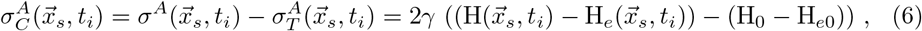

where H_*e*0_ is the average mean curvature of the ellipsoidal mode. Since H_0_ ≃ H_*e*0_ (Fig. 2c), the term H_0_ − H_*e*0_ vanishes in Eq. 6.

**Fig 4.**
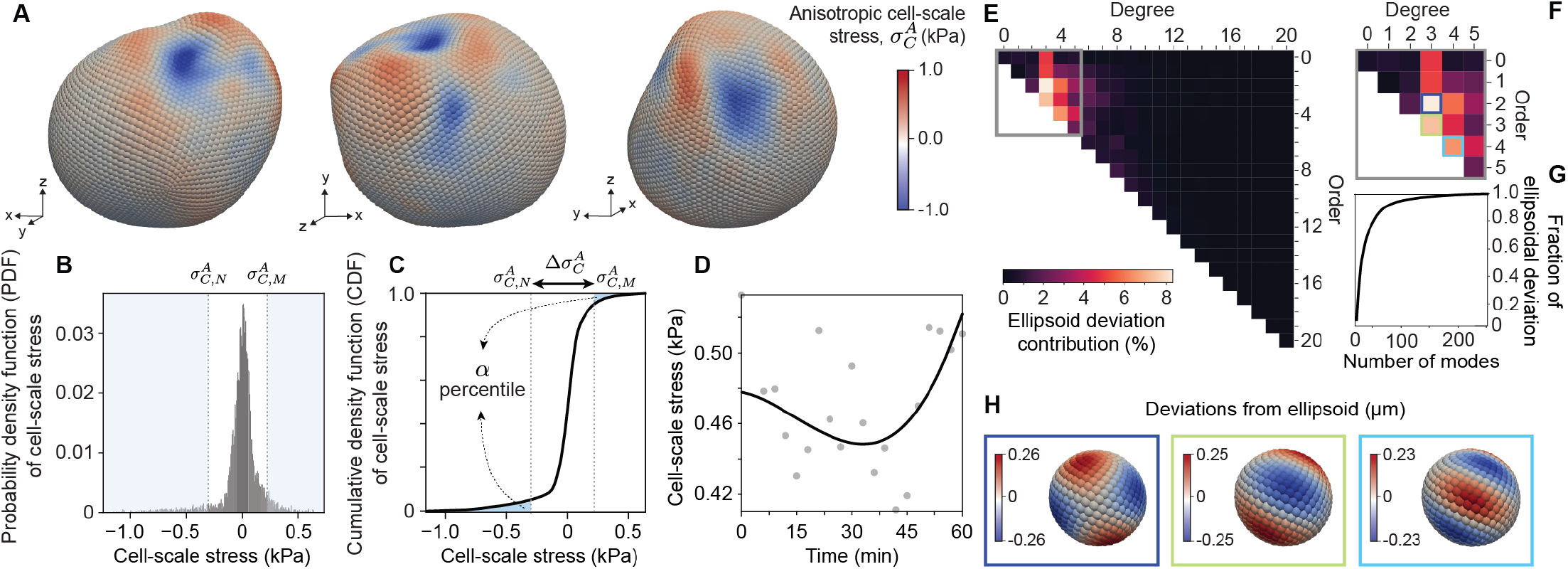
Quantifying cell-scale stresses over time. **A**, Cell-scale stresses shown on the droplet surface from different perspectives at a fixed timepoint. **B-C**, Probability distribution function (or normalized frequency; **B**) and cumulative distribution function (**C**) of cell-scale stresses, 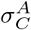, on all 5810 lebedev points. The 5^*th*^ and 95^*th*^ percentile values (*α* = 0.05) define the minimal and maximal cell-scale stresses, 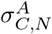 and 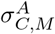 respectively. **D**, Time evolution of the magnitude of cell-scale stresses, 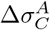. B-spline (black line) shows the trend and gray points are the measured values. **E**, Amplitudes of spherical harmonic modes of the deviations from the ellipsoidal deformation mode. **F**, Highlight from panel *e* showing modes up to degree 5, since these contribute to most of the deviation from the ellipsoid. **G**, Cumulative distribution function of the contribution of each mode to shape deviations from the ellipsoidal mode. The first 50 modes (of over 200) account for 90% of the deviation from the ellipsoidal deformation mode. **H**, Visualization of some of the modes that contribute most to explain the droplet shape deviations from the ellipsoidal mode.

As for the total anisotropic stresses and the tissue-scale stresses, it is important to define a single numerical measure to quantify the time evolution of the magnitude of cell-scale stresses. This can be done in an analogous way as done for the total anisotropic stresses above, by obtaining the normalized distribution of the magnitudes of cell-scale stresses (Fig. 4B), calculating its cumulative distribution (Fig. 4C) and defining maximal and minimal stresses associated with a percentage cut-off, 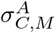 and 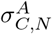, respectively. The time evolution of the magnitude of anisotropic cell-scale stresses is then given by 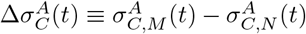 and can be monitored over time (Fig. 4D). The magnitude of cell-scale stresses is larger than that of tissue-scale stresses and is very similar to the measured magnitude of total anisotropic stresses (Fig. 2h), indicating the cell-scale stresses dominate the total magnitude of the stresses.

To better understand the structure of deformation modes higher than the ellipsoidal mode, we decompose the map of cell-scale stresses into spherical harmonics up to degree 20. We use the fact that the coefficients of spherical harmonics show a symmetric structure when representing real quantities and combine the contributions of off-diagonal modes of a given order with their opposite to evaluate the contribution of each mode (Fig. 4E). These combined amplitudes (or coefficients) provide a measure of the relative contribution of each mode to the total droplet deformation. We find that some small number of modes contribute more to the droplet deformations than others (Fig. 4F), but it requires about 50 modes to capture 90% of the droplet deformations (Fig. 4G). The most represented modes (orders 3 and 4) are associated with periodic deformations of the droplet surface at length scales of approximately 12 to 17*μ*m, since the droplet radius is approximately 16*μ*m (Fig. 4H), suggesting the existence of a characteristic length scale of droplet deformations. Beyond the specific example studied here, it is possible to understand the contribution of stresses at different length scales by using mode decomposition.

While deformation modes higher than the ellipsoidal mode provide a better approximation of cellular stresses, they also contain information about deformation modes larger or smaller than the cell size. Beyond specific deformation modes, droplet deformations display maxima and minima at multiple locations on the droplet surface. Since anisotropic stresses compare stresses between different points on the droplet surface, we calculate the anisotropic stresses between pairs of local maxima and minima of stresses. To do so, we first determine the locations of all maxima and minima on the droplet surface by determining the lebedev points with values of the cell-scale stresses larger or smaller to all of their neighbors (Fig. 5A,B; Methods). In order to understand the spatial structure of the extrema on the droplet surface, we calculate the geodesic distances between any pair of maximum and minimum of cell-scale stresses (Fig. 5A; Methods). The distribution of geodesic lengths between them is approximately symmetric and shows that the most frequent distance between all pairs is approximately half the distance between north and south poles of the droplet (Fig. 5C), suggesting equally distributed extrema on the droplet surface. We then evaluate the anisotropic stresses between each pair of local maxima and minima on the droplet, namely the extrema stresses 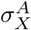, and obtain their distribution (Fig. 5E), which provides information about the spread in the magnitude of anisotropic stresses between extrema on the droplet surface. Our results show a well-defined peak at 80 Pa, but with a distribution characterized by a long tail (average 230 Pa; median 150 Pa). Because all pairs of maxima and minima were considered in this calculation, it is not possible to reveal any local structure associated with these extrema. To characterize the length scales and stresses associated with adjacent extrema (Fig. 5B), we calculate the anisotropic stresses between an extrema (maximum or minimum) and its closest neighbor opposite extrema (Fig. 5D). The distribution of geodesic distances between adjacent pairs of opposed extrema shows a well-defined peak at approximately 3 *μ*m, with the average distance being 4 *μ*m (Fig. 5D), indicating the existence of a well-defined length scale between adjacent extrema at the droplet surface. While the cell diameter is approximately 12 *μ*m in the studied tissues [20], the average length of cell-cell contacts in the tissue is approximately 5 *μ*m [20], indicating that extrema are likely associated with the spatial inhomogeneities in stresses due to cell-cell junctions contacting the droplet surface. Measuring the anisotropic stress distribution of adjacent opposed extrema shows average values of these stresses in the 100-200 Pa range (Fig. 5F).

**Fig 5.**
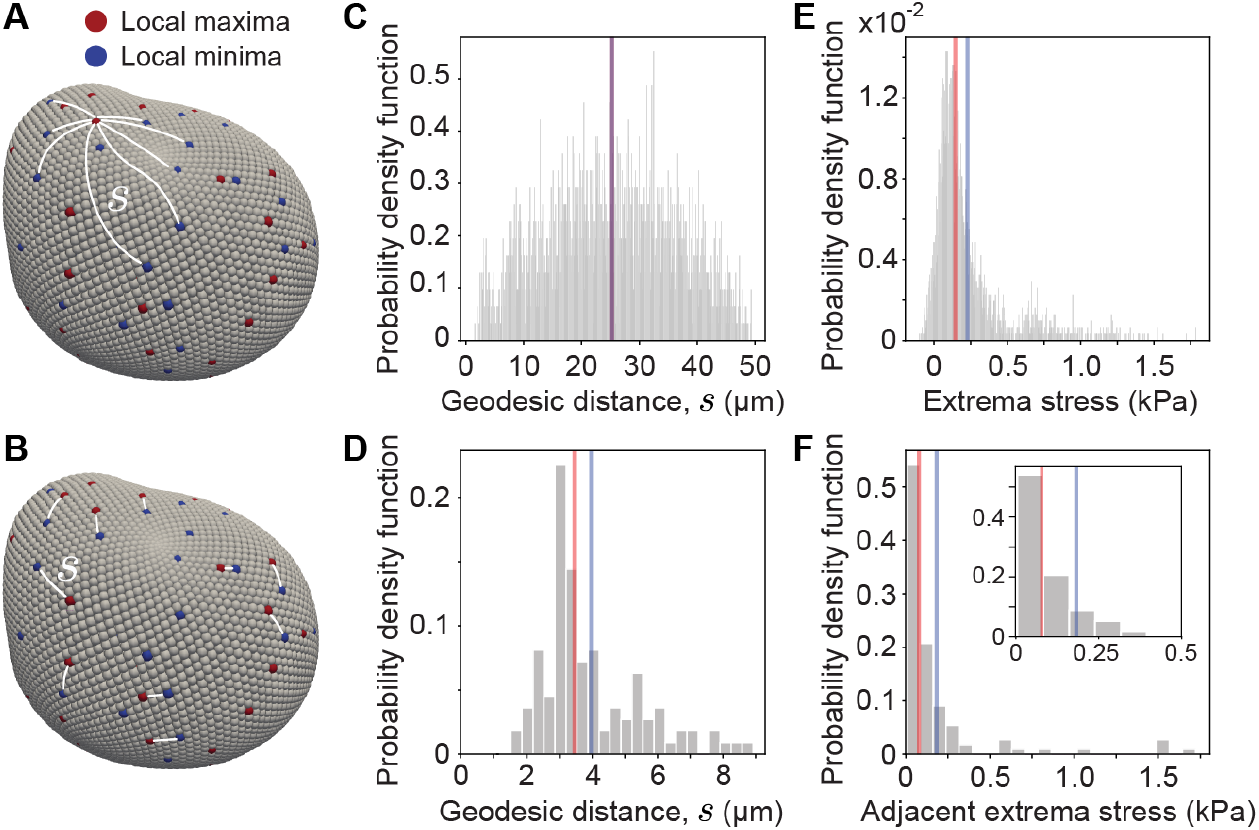
Stresses associated with extrema on the droplet surface. **A-B**, 3D representation of the droplet shape showing the local maxima (red) and minima (blue) on the droplet surface and the geodesic paths (white lines) between a maximum and minima on the surface (**A**) and adjacent pairs of maxima and minima (**B**). **C-D**, Probability density function (normalized frequency) of the geodesic distances between each local maxima and minima (**C**) and between adjacent pairs of maxima and minima (**D**). **E-F**, Probability density function (normalized frequency) of anisotropic stresses between each local maxima and minima (**E**) and between adjacent pairs of maxima and minima (**F**). Mean (blue) and median (red) values are shown for each distribution.

The same analysis of extrema stresses and distances between stress extrema performed here for cell-scale stresses can be done for the total stresses on the droplet surface. Since extrema occur at distances much smaller than the droplet size, the results obtained by considering only cell-scale stresses or the total stresses are very similar.

## Spatial and temporal autocorrelations of anisotropic stresses

The analysis described above enables the characterization of the total, tissue-scale and cell-scale anisotropic stresses and shows the existence of spatial features of the stresses on the droplet. To quantify the degree to which stresses on the droplet are spatially and temporally correlated, we calculate their spatial and temporal autocorrelations, as these provide information about the spatial structure and persistence of stresses, respectively.

Spatial autocorrelations are typically used in condensed matter physics to identify the existence of repetitive spatial structures [33]. In this case, the spatial autocorrelation of anisotropic stresses provides information about the spatial structure of stress inhomogeneities around the droplet. The normalized spatial autocorrelation function is given by

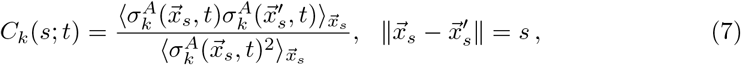

where *s* is the geodesic distance between two given points on the droplet surface, 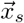 is the coordinate of a point on the surface and *k* indicates that this calculation can be performed for any stress field on the droplet surface (k = total, tissue-scale or cell-scale stresses). Since the spatial autocorrelation is calculated at each timepoint, we obtain the time evolution of the spatial correlations in the system for the total, the cell-scale and the tissue-scale stress anisotropies (Fig. 6A-C). The spatial autocorrelation of total and cell-scale stresses is very similar because the total anisotropic stresses are dominated in magnitude by the cell-scale stresses (Fig. 6A,B). The correlation decays rapidly and reaches a minimum at a length scale of about the cell-size (≃12 *μ*m), displaying a small anti-correlation at this point. In contrast, the variations in the spatial autocorrelation of tissue-scale stresses reflect the geometry of the ellipsoidal mode (Fig. 6C).

**Fig 6.**
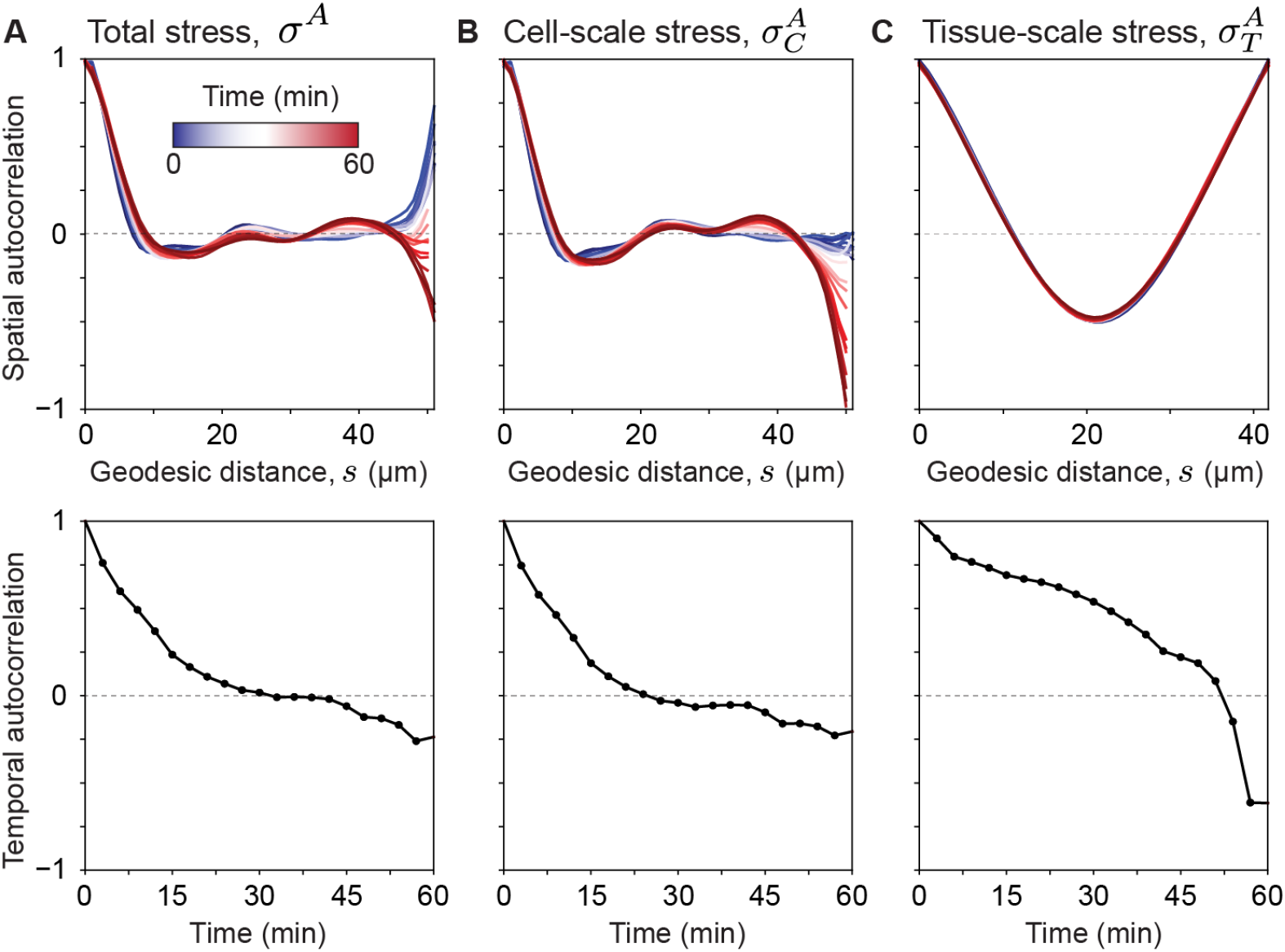
Spatial and temporal autocorrelations of anisotropic stresses. **A-C**, Spatial (top panels) and temporal (bottom panels) auto-correlations of the total anisotropic stresses (**A**), the cell-scale stresses (**B**) and the tissue-scale stresses (**C**).The spatial autocorrelations are shown for each timepoint (color coded), enabling the monitoring of how spatial autocorrelations evolve over time.

Beyond the spatial structure of stress inhomogeneities around the droplet, it is important to characterize how long different stresses persist in the tissue. To do so, we calculate the temporal autocorrelation of anisotropic stresses, which is defined as

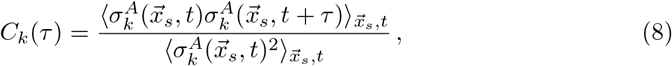

with *τ* being the time difference between the correlated frames. The autocorrelation time, defined as the characteristic timescale over which stresses become uncorrelated, characterizes the temporal persistence of the stresses in the tissue. Cell-scale stresses display the shortest persistence, with stresses becoming uncorrelated at timescales of approximately a few minutes (Fig. 6B). The total stresses show a similar, albeit slightly longer, persistence, in agreement with our results that cell-scale stresses dominate the total stresses (Fig. 6A). In contrast, tissue-scale stresses are much more persistent, with stresses displaying substantial correlations even after 30 minutes, and only change significantly (and sharply) after approximately 50 minutes (Fig. 6C). These results are in agreement with previous analysis done with 2D confocal sections of microdroplets [9], and confirm that tissue-scale stresses are persistent at developmental timescales (approximately 1h).

## Discussion

We have developed STRESS, an automated analysis pipeline to reconstruct the shape of deformable particles and characterize its geometrical features, as well as their time evolution, enabling the quantitative characterization of mechanical stresses in 3D multicellular systems. The STRESS source code, along with installation and usage instructions, can be found at https://github.com/campaslab/STRESS.

The analysis implemented in the STRESS software allows automated surface reconstructions of fluorescently-labeled deformable objects (such as oil microdroplets or gel microbeads) and the analysis of its geometrical features. For the specific case of oil microdroplets, the geometry of the droplet contains the necessary information to quantify mechanical stresses (knowing the droplet interfacial tension). In this case, STRESS allows the automated characterization of the spatiotemporal characteristics of stresses, including the decoupling of cell- and tissue-scale stresses, the measurement of the characteristic length scale of droplet deformations, the analysis of stresses associated to extrema, the spatial correlations of stresses and their time evolution, and the temporal persistence of stresses. Each single droplet thus provides enough statistics to quantify stresses locally in the tissue, from cell to tissue scales and with subcellular resolution. For each analyzed droplet, STRESS automatically generates files for all these quantities (see Supplementary Information or https://github.com/campaslab/STRESS for a user manual).

While using a local representation of the geometry of deformable particles is useful in some cases, especially when it is not possible to reconstruct the entire droplet or a coordinate-free system is more adequate, the global surface representation described herein provides a smoother and more accurate representation of surface deformations that is less sensitive to noise. Moreover, the Gauss-Bonnet test allows a quantification of the error in the reconstruction of the particle deformations, enabling the automated detection of poorly resolved measurements. Finally, the global surface representation allows the calculation of surface integrals, as well as the volume of the droplet with very high accuracy. We believe that accurately calculating the particle volume will be useful for measurements of the isotropic pressure with gel microbeads.

Performing instantaneous measurements of stresses at few timepoints could potentially be done by separately analyzing each timepoint using previous analysis methods. However, the analysis of the time evolution of stresses at high temporal resolution generates a large number of 3D particle reconstructions to analyze, precluding the separated analysis of each time frame. STRESS enables the analysis of the time evolution of stresses from oil microdroplets, *in vivo* and *in situ*, at high temporal resolution.

STRESS automatically decouples the contributions of different deformation modes, enabling the study of stresses associated to different length scales. In the present form, STRESS decouples the ellipsoidal deformation mode, which is associated to tissue-scale stresses (stresses occurring at supracellular scales), and the cell-scale stresses, associated with higher order modes, especially those characterized by length scales similar to the cell size. Moreover, we have developed a new method to characterize stresses associated with extrema (maxima and minima) in mean curvature of the droplet surface, as these extrema are linked to the spatial inhomogeneities in the structure of the tissue (e.g., cellular structure).

Finally, to better understand any spatial features of the stress inhomogeneities in the tissue around the droplet, STRESS calculates the spatial correlations of the measured stresses at each timepoint. This spatial correlation quantifies the length scale over which stresses are correlated on the droplet surface and can be obtained for the stresses associated to any deformation mode. Moreover, the temporal evolution of the spatial autocorrelation provides information about the temporal changes in the spatial inhomogeneities of stresses around the droplet. Finally, for a single droplet, we calculate the temporal autocorrelation of total stresses, as well as tissue- and cell-scale stresses, which provides a measure of how long each of these stresses persist.

Altogether, the STRESS software enables the accurate quantification of multiple types of stresses, as well as their spatial and temporal characteristics, using deformable microdroplets in 3D multicellular systems, from living tissues in developing embryos, to 3D cell culture and organoids.

## Acknowledgments

We would like to thank all members of the Campàs for insightful discussions, and Irene Lim and Ellen Sletten (University of California, Los Angeles) for sharing custom-made fluorinated dyes. Research reported in this publication was supported by the National Institute of General Medical Sciences (award number R01GM135380) and the National Institute of Dental and Craniofacial Research (award number R01DE027620) of the National Institutes of Health, as well as the National Science Foundation (NSF CAREER award, CMMI-1562910). The project was partially supported by the Deutsche Forschungsgemeinschaft (DFG, German Research Foundation) under Germany’s Excellence Strategy – EXC 2068 – 390729961– Cluster of Excellence Physics of Life of TU Dresden.

